# Genetic and environmental factors associated with DNA methylation across childhood and adolescence in individuals with and without attention-deficit/hyperactivity disorder

**DOI:** 10.64898/2026.10.20.760579

**Authors:** Jo Wrigglesworth, Valentine Chirokoff, Peter D. Fransquet, Vicki Anderson, Mark A. Bellgrove, Tim J. Silk

**Affiliations:** Deakin Lifespan Institute and School of Psychology, Deakin University, Burwood, VIC, Australia; Institute for Mental and Physical Health and Clinical Translation (IMPACT), School of Medicine, Deakin University, Geelong, VIC, Australia; Public Health Genomics, School of Public Health and Preventive Medicine, Monash University, Melbourne, VIC, Australia; Murdoch Children’s Research Institute, Melbourne, VIC, Australia; Department of Psychology, Melbourne School of Psychological Sciences, The University of Melbourne, VIC, Australia; Turner Institute for Brain and Mental Health, School of Psychological Sciences, Monash University, Clayton, VIC, Australia

**Author notes:** Correspondence: Dr Jo Wrigglesworth, School of Psychology, Deakin University, Burwood, Australia. Jo Wrigglesworth and Valentine Chirokoff are joint first authors.

## Abstract

**Background:** Genetic and environmental factors both contribute risk to the development of childhood Attention-deficit/hyperactivity disorder (ADHD), yet the molecular mechanisms conferring this risk remain unclear. We investigated the genetic and environmental factors influencing DNA methylation (DNAm) at sites linked to ADHD symptom trajectories.

**Methods:** We used two overlapping subsamples from a longitudinal paediatric cohort: (1) children with genetic and epigenetic data at first wave saliva sampling (Wave 3, n=113, 9.47-11.86 years) underwent methylation quantitative trait loci (meQTL) analysis; (2) children with environmental (Waves 1-3) and one or more waves of epigenetic data (n=126, 9.5-11.9 years) were analysed using LASSO regression to identify predictors of ADHD (Remitted ADHD/Persistent ADHD/Controls) and Wave 3 DNAm. In a subset with multiple DNAm timepoints (n=75), linear mixed models tested whether environmental predictors associated with longitudinal DNAm change, and whether ADHD moderated these associations. Analyses were conducted on 12 CpGs in which we previously identified change in DNAm according to the ADHD clinical outcome.

**Results:** Nineteen meQTLs for 5 CpGs explained up to 40% of DNAm variance. We identified 17 environmental predictors of ADHD and wave 3 DNAm. Higher household income and school quality of life (QoL) were associated with reduced DNAm at cg02208776 (household) and cg11124426 (QoL) over time. Critically, higher income in Remitted ADHD showed greater DNAm declines than Controls, with opposite patterns in Persistent ADHD.

**Conclusion:** DNAm patterns linked to ADHD outcomes were associated with genetic and environmental factors. Notably, household income and school functioning were associated with epigenetic change, with income effects varying by ADHD trajectories - characterising higher income as protective in remittent individuals.

## INTRODUCTION

Attention-deficit/hyperactivity disorder (ADHD) is a prevalent neurodevelopmental condition, impacting approximately 5% of children and adolescents worldwide (Cortese et al., 2023). There is considerable heterogeneity in symptom profiles between individuals, which can also fluctuate within an individual throughout the course of development (Franke et al., 2018; Sibley et al., 2022). For some children with ADHD, their symptoms remit (reduced to subthreshold levels) by the time they reach young adulthood, while for others the symptoms remain chronic (Sibley et al., 2022; Van Meter et al., 2024). Recent evidence-based consensus suggests that both genetic and environmental factors influence the emergence and course of ADHD (Faraone et al., 2021).

It is evident that there is a significant genetic component to ADHD, with heritability estimates as high as 74% in twins (Faraone & Larsson, 2019). However, genome-wide studies revealed how only a small proportion of genetic risk for ADHD (22%, Demontis et al., 2019) can be attributed to common genetic variants (variation found in more than 1% of the population, Faraone & Larsson, 2019). At the same time, ADHD has been linked to several environmental factors (Faraone et al., 2021). More recently, there has been growing interest in epigenetics, which might provide a mechanistic account of how environmental factors may play a role in regulating gene expression to influence health outcomes. The most widely studied epigenetic mechanism in ADHD is DNAm. DNAm can regulate gene expression in response to genetic and environmental stimuli through the addition of a methyl molecule to a cytosine of a cytosine-phosphate-guanine ‘CpG’ dinucleotide (Jaenisch & Bird, 2003). Interest stems from evidence of DNAm playing a regulatory role in brain development (Gapp et al., 2014) and the association with other neurodevelopmental disorders (Dall’Aglio et al., 2018). Moreover, the ease of collecting and evaluating DNAm from peripheral tissue is especially useful when studying paediatric populations (Cecil & Nigg, 2022). Several cross-sectional epigenome-wide studies (EWAS) have identified CpGs where DNAm differed between children and adolescents with and without ADHD (e.g., Mooney et al., 2020; Walton et al., 2017; Wilmot et al., 2016). Longitudinally, our own work has identified 12 CpGs where change in DNAm varied according to the clinical outcome of ADHD remittance versus persistence (Wrigglesworth et al., 2026).

DNAm trajectories are shaped by genetic and environmental influences. Genetic influences on DNAm variation appear to remain relatively stable from childhood onwards, whereas the influence of environmental exposures on DNAm variation increases across the lifespan (Gaunt et al., 2016). Among individuals with ADHD, epigenetic studies have identified methylation quantitative trait loci (meQTL, single nucleotide polymorphisms ‘SNPs’ associated with DNAm at nearby or far away sites) which suggest a genetic contribution on differences in DNAm (Barker et al., 2018; Goodman et al., 2020; Mooney et al., 2020). Studies have also highlighted several early life and childhood risk factors for ADHD that are associated with DNAm, particularly those during the prenatal period (e.g., Breton et al., 2016; Sharp et al., 2017). The most robust finding relates to maternal smoking, where long-term perturbations in DNAm have been linked to prenatal exposures (Joubert et al., 2016; Kupers et al., 2015; Lee et al., 2015; Richmond et al., 2015; Rzehak et al., 2016; Toledo-Rodriguez et al., 2010). The biological embedding of adverse childhood experiences is also widely studied through epigenetics, with findings linking ACE-related DNAm patterns with mental health outcomes (Nothling et al., 2020; Parade et al., 2021). While prenatal environments have been examined by ADHD-related DNAm studies (Chaumette et al., 2023; Choi et al., 2023; Frey et al., 2018; Gervin et al., 2017; Olstad, Nordeng, Lyle, et al., 2023; Olstad, Nordeng, Sandve, et al., 2023), investigations of childhood risk factors are lacking. A comprehensive epigenetic analysis incorporating both periods, and examining variation across clinical trajectories is needed.

Extending our prior work (Wrigglesworth et al., 2026), this study aimed to determine whether 1) genetic variants (i.e., meQTLs) are cross-sectionally associated with late childhood DNAm; 2) early life and childhood predictors of ADHD status (Remitted ADHD/Persistent ADHD/Controls) are cross-sectionally associated with DNAm in late childhood, and 3) clinically relevant early life and childhood environmental predictors of late childhood DNAm are also associated with intra-individual (within-person) change in DNAm across adolescence, and whether this varies according to ADHD status.

## METHODS

### Study participants

Data were derived from the larger longitudinal Children’s Attention Project (CAP, Sciberras et al., 2013) and its ancillary neuroimaging study, the Neuroimaging of the Children’s Attention Project (NICAP, Silk et al., 2016). Briefly, a community sample of children (aged 6–8 years) were initially recruited from 43 socioeconomically diverse primary schools in Melbourne, Australia, with follow-up for up to five timepoints with 18-month intervals. Participants were screened for ADHD using independent parent and teacher reports on the Conners 3 ADHD index (Conners, 2008), before receiving a confirmed ADHD diagnoses using the NIMH Diagnostic Interview Schedule for Children-IV (DISC-IV, Shaffer et al., 2000). Diagnosis was reassessed at the third and fifth wave follow-up. For this study, participants were classified as meeting criteria for including in the ADHD group if they met diagnostic criteria at the initial recruitment or wave 3 follow-up. The ADHD group was then further parsed into those who continued to meet diagnostic criteria at their last assessment (Persistent ADHD), and those who no longer met criteria by their last assessment (Remitted ADHD). Control participants did not meet criteria for ADHD at any wave.

At each wave, multiple direct assessments were performed, and parent- and teacher-reported questionnaires were collected, providing measures pertaining to several potential protective and risk factors including individual, health, and retrospective prenatal, school, home and life stress factors (See **Supplementary Table 1** for details). Across waves 3-5, a subset of participants also contributed saliva samples for epigenetic and genetic analysis.

The current study included participants who met at least one of the following criteria (**Figure 1**): 1) complete genotypic and epigenetic data at the first wave of biological sampling (Wave 3=113); 1) available historical environmental data (Waves 1-3) plus epigenetic data collected at one or more waves (n=126). The study protocol, including all assessment procedures, was approved by the ethical committees of The Royal Children’s Hospital and Deakin University (Melbourne, Australia), and written informed consent was obtained from the parents or guardians of all participating children.

**Figure 1.**
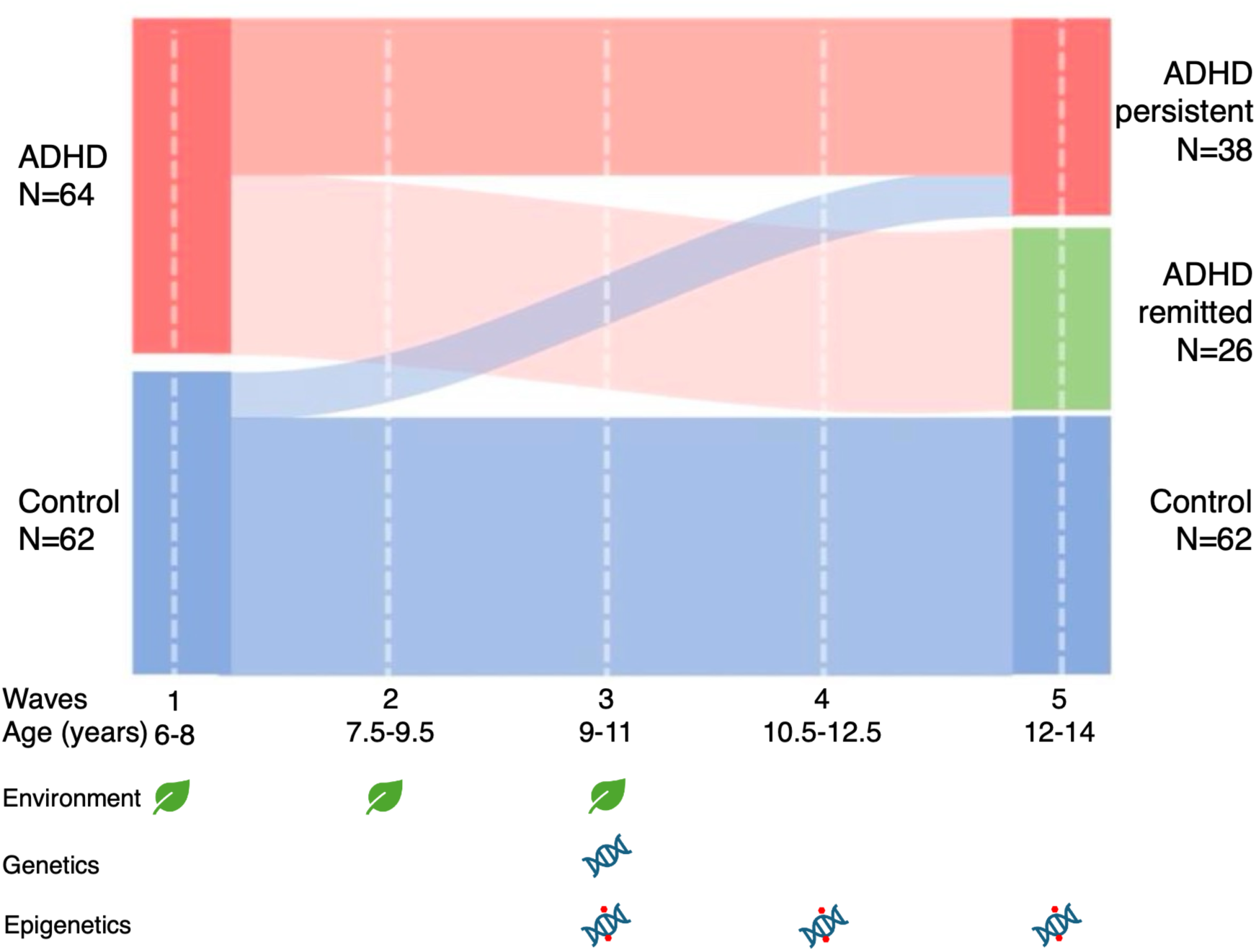
Summary of study design and data collection. Legend: ADHD frequency based on sample of participants with available environmental data (Waves 1-3) and epigenetic data at one or more waves.

### Genetic variants

DNA extracted from saliva was genotyped using the Illumina Global Screening Array (GSA-MD-24v1- 0; Illumina, San Diego, CA, USA) and initially pre-processed at the Stanley Center for Psychiatric Disease, Broad Institute (Cambridge, MA, USA). A series of quality control (QC) steps were performed, and are detailed in the **Supplementary Methods**. Briefly, initial QC procedures were executed using the Ricopili pipeline (Lam et al., 2020), which removed low-quality samples and variants after three consecutive runs. Genetic ancestries were inferred using the 1000 Genomes Project reference panel (1000 Genomes Project Consortium et al., 2015), and samples clustering within the European ancestry were retained for downstream analyses.

Missing genetic data were imputed using the TOPMed Imputation Server (https://imputation.biodatacatalyst.nhlbi.nih.gov) with the NHLBI TOPMed Reference Panel (Version r3, GRCh38). Low-quality variants were removed during post-imputation QC either if the imputation quality score R² ≤ 0.3 or the minor allele frequency (MAF) ≤ 0.05. A total of 8,231,947 variants were retained for downstream analyses, with 96.02% of the input variants overlapping the reference panel.

### Environmental exposures

The World Federation of ADHD summarised evidence on the environmental correlates of ADHD risk in an international consensus statement in 2021 (Faraone et al., 2021). We used this knowledge to focus our epigenetic analyses on environmental factors related to ADHD. For interpretation purposes, we grouped exposures according to the ‘*Child’s health*’, ‘*Home environment*’, ‘*Perinatal*’, ‘*School environment*’ and ‘*Stressful life events*’, and provide a list of the analysed variables in **Supplementary Table 1**.

Environmental data on the child’s overall health, sleep habits, medication use, perinatal factors, bullying exposure, parental mental health, and socio-economic characteristics were collected using a combination of standardised scales and study-designed questionnaires. Assessments include the Conners’ 3 ADHD index (Conners, 2008), the Spence Children’s Anxiety Scale (Spence et al., 2014), the Social Experience Questionnaires (SEQ, Crick & Grotpeter, 1996), the Paediatric Quality of Life Inventory (PedsQL, Varni et al., 2003), the Conners’ Adult ADHD Rating Scale (CAARS, Conners et al., 1999), the Kessler Psychological Distress Scale (K6, Furukawa et al., 2003), the five positive affect items on alertness, inspiration, determination, attentiveness and activeness from the Positive and Negative Affect Schedule Short Form (PANAS-SF, Watson et al., 1988), the Longitudinal Study of Australian Children (LSAC) Parenting Scale (Wake et al., 2014), and the Stressful Life Events Questionnaire (SLEQ, Brugha & Cragg, 1990).

Ground-level ambient air pollution exposure was quantified for fine particulate matter (PM2.5) and Nitrogen Dioxide (NO2) by sourcing openly shared datasets, made available by NASA’s Earth Science Data and Information System (ESDIS) Project. This includes the Global Annual PM2.5 average (Battelle Memorial Institute; Center For International Earth Science Information Network-CIESIN-Columbia University, 2013) and the three-year average NO2 grids (Geddes et al., 2017). Using QGIS software (version 3.40.11, https://github.com/qgis/QGIS), the median exposure levels from PM2.5 and NO2 grids, at a time that aligns with the preceding year of epigenetic assessment, were extracted for the postcode associated with the participant’s home residence in the Melbourne region.

The current study derived ‘*Absolute*’ (coded ‘1’ to perinatal factors, parental mental health history, child medication history, and all stressful life event exposures if ‘Yes’ at any timepoint) and ‘*Overall*’ (applied to continuous exposures, representing median across Waves 1-3) measures of environmental exposure using data collected across Waves 1-3. Measures are the final output after coding and QC procedures were performed, which we describe in the **Supplementary Methods**.

### DNA methylation preprocessing and quality controls (QC)

DNA from saliva was bisulphite treated and hybridised to the Infinium Human Methylation 850 (EPIC) BeadChip arrays, version 1.0 (Illumina, San Diego, CA, USA) at NTX-Dx (Diagenode, Ghent, Belgium). DNAm data underwent QC procedures which follow the Maksimovic and colleagues (2016) Bioconductor workflow (Maksimovic et al., 2016), using the R *minfi* package (Aryee et al., 2014). Details regarding QC are described elsewhere (Wrigglesworth et al., 2026). Briefly, data across all visits were normalised collectively using subset quantile normalisation (Touleimat & Tost, 2012; Wu & Aryee, 2010). Probes were removed if they had a poor-quality signal (113,011 with detection p>0.01), were located on a sex chromosome (n=16,376), overlapped a SNP at the CpG or extension site (n=26,455), or were cross-hybridising (n=34,471 mapped to multiple locations) (Pidsley et al., 2016). Probes were originally mapped to genomic features of the human genome assembly GRCh37 (hg19) using *IlluminaHumanMethylationEPICanno.ilm10b4.hg19.* For the current study, after preprocessing was complete, probes were lift-over to the GRCh38 (hg38) genome assembly. M-values, representing the ratio of methylated/unmethylated intensity, were extracted from the entire epigenome (n=675,546) and used in downstream analyses (Du et al., 2010). Extending on our previous study findings (Wrigglesworth et al., 2026), we analysed 12 CpGs where longitudinal patterns of DNAm differed between ADHD groups.

Cell-type heterogeneity on DNAm was controlled using the EpiDISH ‘*centEpiFibIC.m*’ reference panel (Zheng et al., 2020). This panel has been shown to be appropriate when estimating epithelial and immune cells from paediatric saliva samples (Chan et al., 2026). Given that these cell types are inversely proportional, we only included epithelial proportion as a covariate in downstream analyses, which is mathematically equivalent to adjusting for leukocyte proportion.

## Statistical analyses

The aims of determining whether genetic and environmental factors are associated with DNAm were addressed using two independent approaches: 1) focuses on the role of genetic variation through meQTL analyses; 2) examines whether early life and childhood environmental predictors of ADHD status were cross-sectionally associated with wave 3 DNAm, and whether those significant predictors were associated with longitudinal intra-individual change in DNAm. We summarise these two approaches in the subsections below.

### 1) Methylation quantitative trait loci (meQTL) analyses

This study used the *Matrix eQTL* R package (v2.3, Shabalin, 2012) to test both local ‘*cis*’ (SNPs within ±1Mb of a CpG site) and distant ‘*trans*’ (SNPs beyond 1Mb, including inter-chromosomal) genetic effects on wave 3 DNAm, adjusting for age, sex, ADHD status, batch (processing plate), epithelial cell-proportion and population stratification (first 10 PCs of the genetic data). Multiple correction was applied separately to each analysis using the Benjamini-Hochberg false discovery rate (BH-FDR, Benjamini & Hochberg, 1995).

A custom greedy LD-clumping algorithm was then used to reduce FDR-significant associations at each CpG to independent lead SNPs. This custom algorithm follows a similar logic to PLINK, which we described in detail in the **Supplementary Methods**. Each lead SNP’s effect was characterised by its regression beta coefficient (estimated change in DNAm M-value per additional copy of the effect allele under the additive model) and partial R^2^ (unique variance in DNAm explained beyond the covariates).

### 2) Environmental predictors of DNA methylation

To investigate early life and childhood environmental predictors of ADHD-related DNAm, we conducted a series of cross-sectional and longitudinal analyses.

*Cross-sectionally*, we identified clinically relevant early life and childhood environmental predictors of DNAm using a two-step approach. This approach sequentially applied Least Absolute Shrinkage and Selection Operator (LASSO) regression models: the first step identified nominally significant environmental predictors of ADHD status, and the second step determined whether these environments are also associated with wave 3 DNAm. This two-step approach is detailed in **Supplementary Methods**.

*Longitudinally*, we incorporated our cross-sectional findings to determine whether clinically relevant environmental predictors of wave 3 DNAm were also associated with intra-individual change in DNAm. We further examined whether associations were moderated by ADHD status.

The identified environmental factors were used as predictors in linear mixed models (LMM), first testing longitudinal associations with intra-individual change (within-person) in DNAm across the whole cohort, using the *lmerTest* package (Kuznetsova et al., 2017). LMMs were run separately for each environmental predictor and its associated CpG, along with the interaction between environment and time. Time represents a person-mean centred change, calculated as the difference between each participant’s chronological age at each wave and their age at the first assessment (age at any wave – age at first wave). All models were adjusted for ADHD (Remitted ADHD/Persistent ADHD vs. Controls), sex, epithelial cell-proportion, batch (i.e., plate), and medication use (unless main predictor was medication use); each participant was included as a random intercept.

The moderating effect of ADHD status on the relationship between environment and intra-individual change in DNAm was examined using the same adjusted LMM, but included ADHD status as a three-way interaction (ADHD status * environment * time) rather than as a covariate. We performed two sets of group comparisons: 1) ADHD groups (Remitted or Persistent ADHD) vs. Controls, and 2) Remitted ADHD vs. Persistent ADHD. FDR-significant associations were further interpreted within each diagnostic group (Remitted ADHD, Persistent ADHD and Controls) using adjusted LMM, stratified by follow-up diagnosis.

Across all analyses, we removed models which were non-converging due to insufficient random-effects variance across participants. For each remaining CpG model, p-values for environmental predictors were corrected for multiple testing using the BH-FDR (Benjamini & Hochberg, 1995), with a significance set at FDR < 5%. Continuous predictors were standardised, and binary categorical variables were centred (using -0.5, and 0.5 values). Diagnostic plots visually confirmed that each model fitted data from our smaller subsample of participants with repeated biological samples.

## RESULTS

### Genetic variation influencing childhood DNA methylation

We examined local (*cis*) and distant (*trans*) genetic effects on Wave 3 DNAm using data from 113 children (mean age of 10.37 years, ranging 9.47 to 11.86 years, 63% were male). Among these participants, 55 had a diagnosis of ADHD at recruitment.

Across all analyses, we identified 19 independent meQTL signals (2 cis and 17 trans), which involved 5 of the 12 CpG sites (cg02208776, cg12881363, cg17601216, cg17657037, cg21443143). All 19 lead SNPs showed large partial R² values (19.3–40.9%), indicating that individual SNP genotypes explained a substantial proportion of inter-individual DNAm variance at these loci after adjustment for covariates. SNPs implicated both protein-coding genes (18/19 nearest genes) and a single lncRNA (*LINC01440*), with all annotated protein-coding genes spanning diverse biological functions including RNA metabolism (*MOV10L1*), DNA repair (*MGMT*), glycosylation (*ST3GAL1*, *B3GLCT*), lipid metabolism (*ELOVL6*, *NUDT12*), and ion channel function (*CACNA1D*). Seven of the 12 CpGs (cg15550572, cg02968116, cg11124426, cg12577151, cg19892195, cg00060320, cg26901352) showed no per-CpG FDR-significant meQTL associations in either cis or trans. Each analysis results are described in subsequent sections.

### Cis-meQTL Associations

At per-CpG FDR-significance, 189 cis associations were identified across two of the 12 CpGs (cg12881363 and cg21443143). No cis-meQTL signals were identified for the remaining 10 candidate CpGs. LD clumping (r² ≥ 0.10, ±500 kb) collapsed 189 associations into two independent cis signals (see **Figure 2**).

**Figure 2.**
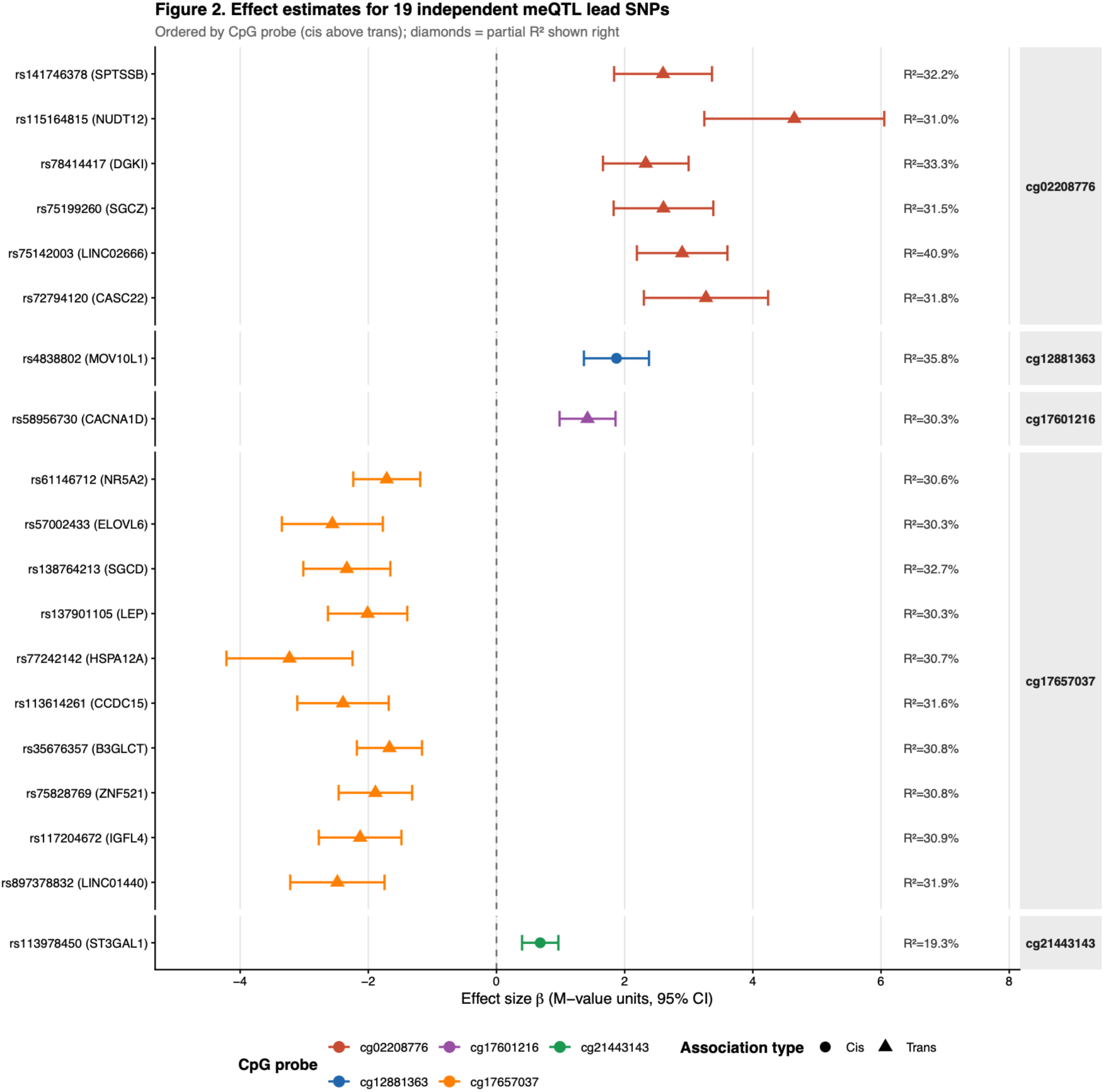
Results from the cis- and trans-meQTL analyses, after LD clumping, representing the effect size (ß), and 95% confidence interval of the SNP effects on CpG DNA methylation (DNAm). *Legend: Direction of associations between SNP variants and DNAm vary per sites with negative effects size representing an opposing link. Sites represented by points highlight SNPs underlying DNAm at a nearby site (±1Mb from CpG, cis-meQTL), whereas triangles represent genetic influences on DNAm at distant sites (>1Mb from CpG, including inter-chromosomal, trans-meQTL). R^2^ represents DNAm variance explained per SNP sites, and range from 19.3 to 40.9% indicating a significant genetic influence on DNAm*.

cg12881363: the lead SNP rs4838802 (chr22:50,109,283) showed the strongest cis-meQTL association (β = 1.872 M-value units, 95% CI: 1.364–2.380; p = 7.67 × 10⁻¹¹), explaining 35.8% of DNAm variance at cg12881363, after adjustment. This signal encompassed a clump of 188 SNPs in strong LD spanning the *MOV10L1* locus on chromosome 22.

cg21443143: rs113978450 (chr8:133,501,650) was the sole (lead) SNP significantly associated with cg21443143 in cis (β = 0.682, 95% CI: 0.399–0.966; p = 6.20 × 10⁻⁶; partial R² = 19.3%). The SNP resides within the *ST3GAL1* gene body on chromosome 8.

### Trans-meQTL Associations

At per-CpG FDR-significance, 31 trans associations were identified across three of the 12 CpGs (cg02208776, cg17657037 and cg17601216). LD clumping identified 17 independent trans signals (**Figure 2**).

cg02208776: the strongest trans association in the entire analysis was rs75142003 (chr10:129,796,690, nearest gene: *MGMT*; β = 2.897, 95% CI: 2.191–3.603; p = 1.41 × 10⁻¹²; partial R² = 40.9). Five additional independent trans signals for cg02208776 were identified on chromosomes 3 (*SPTSSB*, rs141746378), 5 (*NUDT12*, rs115164815), 7 (*DGKI*, rs78414417), 8 (*SGCZ*, rs75199260), and 16 (*TOX3*, rs72794120), each explaining 31.0–33.3% of DNAm variance.

cg17657037: ten genome-wide independent signals were identified for cg17657037, with lead SNPs on chromosomes 1, 4, 5, 7, 10, 11, 13, 18, 19, and 20. Effect directions were uniformly negative (β range: −1.671 to −3.231), and partial R² values were consistent across signals (30.3–32.7%). Implicated genes include *SGCD*, *LINC01440*, *ZNF521*, *B3GLCT*, *HSPA12A*, *NR5A2*, *ELOVL6*, *LEP*, *IGFL4*, and *CCDC15*. cg17601216: a single trans signal was identified for cg17601216 - rs58956730 (chr3:53,809,542, *CACNA1D*; β = 1.421, 95% CI: 0.984–1.857; p = 4.31 × 10⁻⁹; partial R² = 30.3%).

### Environmental predictors of DNA methylation

We determined whether clinically relevant early life and childhood environments are cross-sectionally associated with DNAm (‘*Cross-sectional DNA methylation in late childhood*’), and longitudinally associated with intra-individual change in DNAm across the whole cohort (‘*Longitudinal change in DNA methylation*’). Longitudinal investigations include additional analyses to test for moderation by ADHD status.

### Cross-sectional DNA methylation in late childhood

We cross-sectionally examined clinically relevant early life and childhood environmental predictors of wave 3 DNAm. Analyses involved 126 children (ages ranging between 9.5 to 11.9 years, 64% were male) with available environmental data and Wave 3 epigenetic data (63% were also eligible for meQTL analyses). We characterise this sample using all the surviving QC environmental factors in **Supplementary Table 2**. Of these participants, 64 had a history of ADHD, of which 59% continued to meet criteria for ADHD at their last wave (Persistent ADHD); the remaining 41% no longer met criteria (Remitted ADHD).

Across two steps of analysis, we applied LASSO models to identify environmental predictors associated with ADHD status (Step 1), and that are also associated with Wave 3 DNAm (Step 2). We describe the findings in the **Supplementary Results**. Briefly, 23 environmental factors were selected as predictors of ADHD status: 14 predicted Remitted ADHD vs. Controls; 10 predicted Persistent ADHD vs. Controls, and 6 predicted Remitted ADHD to Persistent ADHD (**Figure 3**). These predictors covered a range of environmental categories, including Air pollution, Home environment, School environment, Perinatal, and Stressful life events, with most relating to the Child’s health (n=12). Among the environmental predictors of ADHD status, 17 were associated with DNAm at 6 CpG sites (**Figure 3**) - cg02208776 (*HSPE1;HSPD1)*, cg02968116, cg11124426, cg12577151 (*ACAD9*), cg1281363 (*MOV10L1)* and cg17657037 (*ITGB1BP3)*.

**Figure 3.**
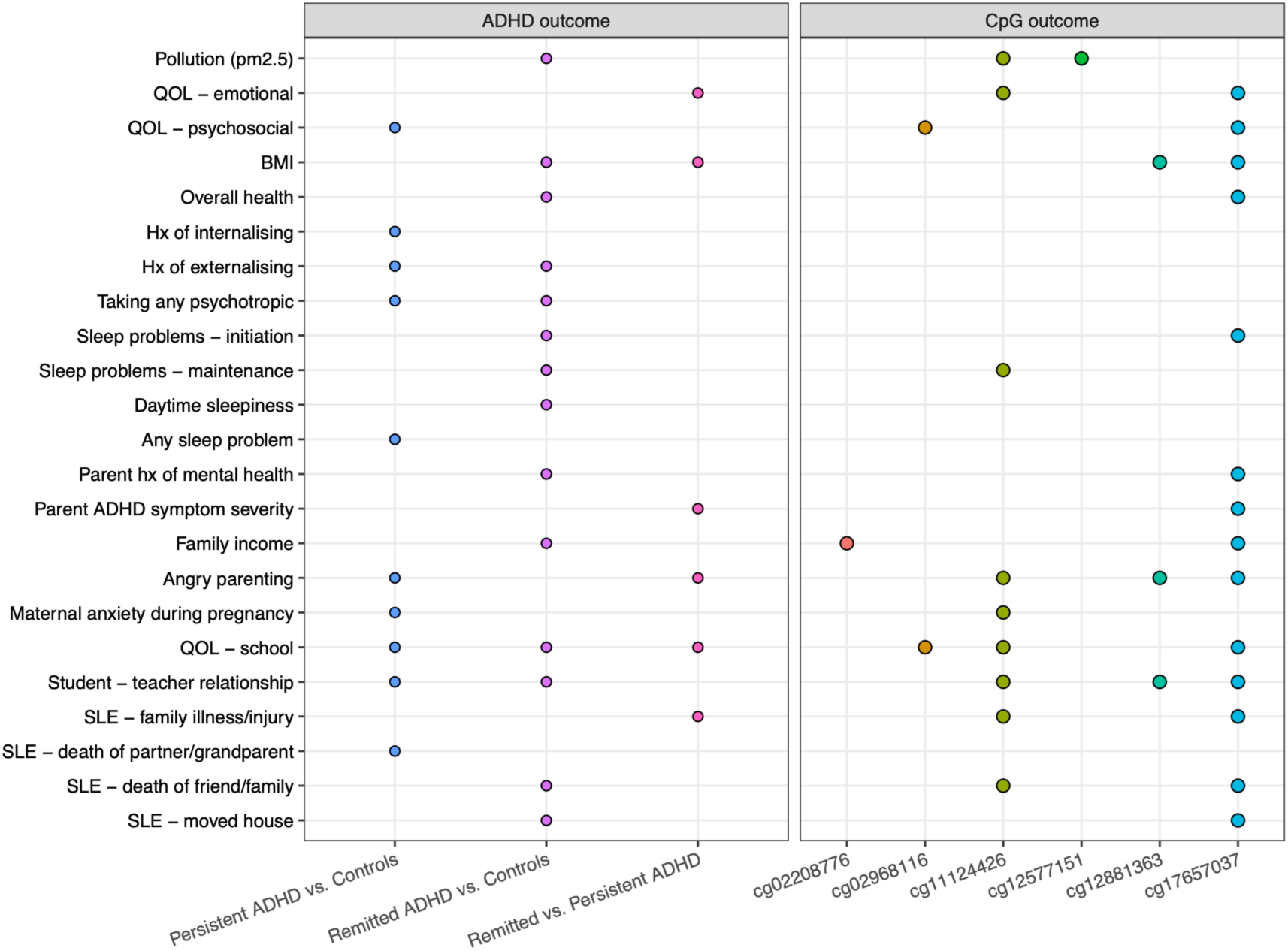
LASSO model selection of environmental factors predicting three pairwise ADHD group comparisons (‘ADHD outcome’) and first wave DNA methylation at 12 CpG sites (‘CpG outcome’). *Models were adjusted for child age and sex. BMI=Body Mass Index; QoL=Quality of Life; SLE=Stressful life event*.

### Longitudinal change in DNA methylation

The set of clinically relevant early life and childhood environmental predictors of Wave 3 DNAm were further examined for longitudinal associations with intra-individual change in DNAm - first across the whole cohort, and then by ADHD status. To conduct these analyses, we reduced the larger sample of participants with environmental data (n=126) to 75 with epigenetic data across two (n = 46) or three (n = 29) waves (mean age of 10.4 ± 0.43 years, 69% were male). Among this sample, 34 participants had a childhood history of ADHD, of which 19 continued to meet diagnostic criteria at their final wave (Persistent ADHD). Across each diagnostic group, there was a greater representation of males than females. The environmental predictors used in this longitudinal investigation are summarised in **Supplementary Table 5**. Fifty-one participants were not eligible for this portion of study as they had only provided a single biological sample. Similarities and differences between these 51 individuals and the eligible participants for longitudinal analyses in the **Supplementary Results**.

Across the whole cohort we examined whether there were early life and childhood environmental predictors of intra-individual change in DNAm, independent to ADHD status (**Table 1**). Among these results were two CpGs (cg02208776 [*HSPE1;HSPD1*], cg11124426) with findings that survived FDR correction: higher overall household income was associated with intra-individual declines in cg02208776 DNAm, and higher overall score of school QoL was associated with intra-individual declines in cg11124426 DNAm (**Figure 4**).

**Figure 4.**
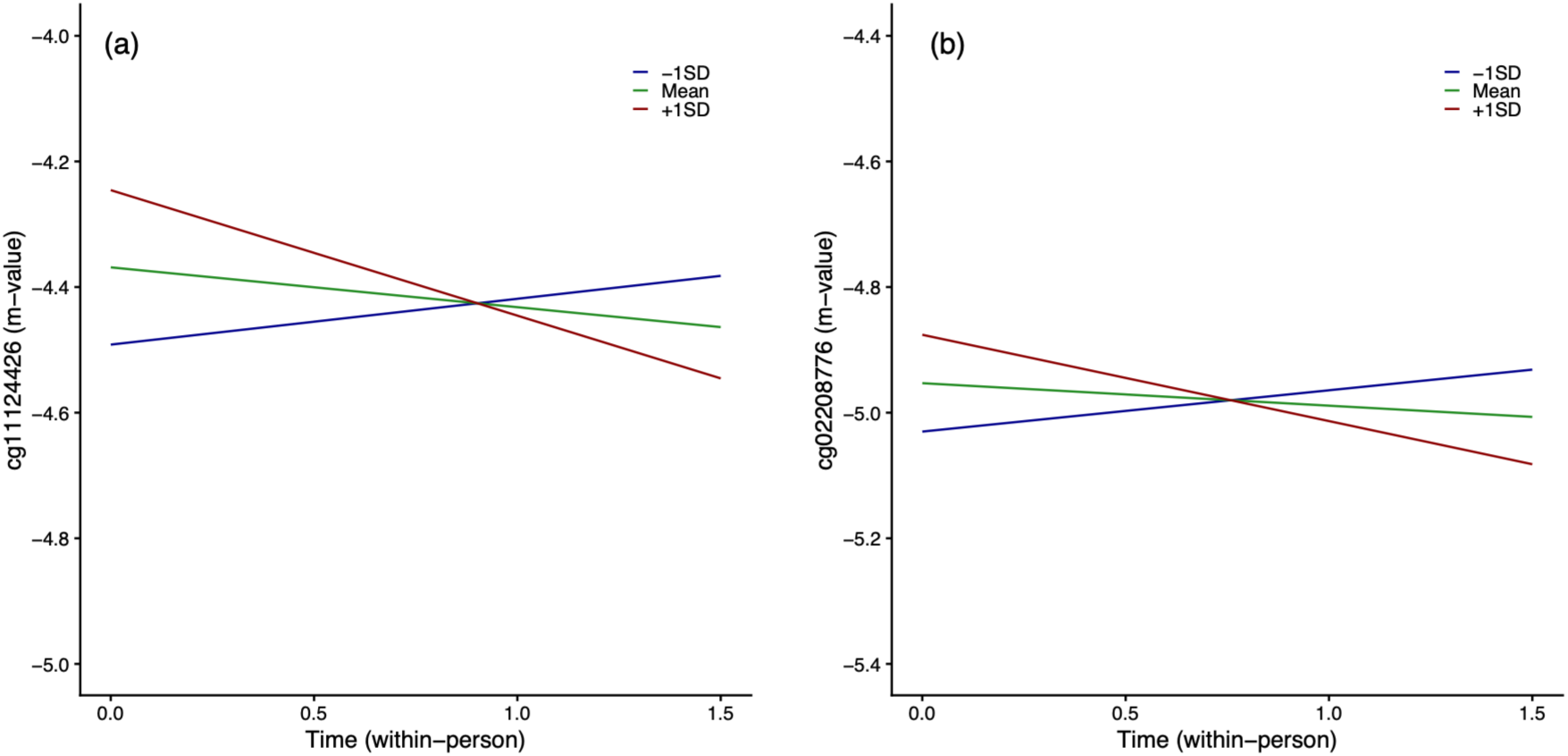
Results from linear mixed models examining associations between (a) school quality of life (QoL) or (b) household income and intra-individual change in DNA methylation (DNAm) across the whole cohort. *Legend: Time represents a person-mean centred change, calculated as the difference between each participant’s chronological age at each wave and their age at the first assessment (age at any wave – age at first wave), which was used to model within-person change in DNAm*.

**Table 1.** Results from linear mixed models testing associations between prenatal and childhood environmental exposures and individual change in DNA methylation in the total study sample.

| Exposure | Probes | Estimate (95% CI) | p |
| --- | --- | --- | --- |
| **Child's health:** |  |  |  |
| QoL: emotional functioning | cg11124426 | 0.04(-0.06, 0.15) | 0.415 |
|  | cg17657037 | -0.09(-0.19, 0.02) | 0.109 |
| QoL: school functioning | cg11124426 | -0.14(-0.23, -0.04) | **0.006** |
|  | cg17657037 | 0.02(-0.07, 0.12) | 0.629 |
| QoL: psychosocial health summary school | cg17657037 | -0.003(-0.10, 0.10) | 0.948 |
| BMI | cg12881363 | -0.003(-0.13, 0.12) | 0.963 |
|  | cg17657037 | -0.05(-0.15, 0.04) | 0.294 |
| Parents rating of child's overall health | cg17657037 | 0.09(-0.01, 0.19) | 0.083 |
| **Home environment:** |  |  |  |
| Parent ADHD symptom severity | cg17657037 | 0.02(-0.09, 0.13) | 0.743 |
| Household income | cg02208776 | -0.10(-0.18, -0.02) | **0.018** |
|  | cg17657037 | -0.03(-0.13, 0.07) | 0.579 |
| Angry parenting | cg11124426 | -0.002(-0.11, 0.10) | 0.974 |
|  | cg12881363 | -0.02(-0.15, 0.11) | 0.731 |
|  | cg17657037 | 0.02(-0.08, 0.13) | 0.704 |
| **Pollution:** |  |  |  |
| Air particulate matter (pm) 2.5 | cg11124426 | 0.001(-0.10, 0.10) | 0.988 |
|  | cg12577151 | 0.01(-0.04, 0.06) | 0.717 |
| **School environment:** |  |  |  |
| Student teacher relationship - total | cg11124426 | -0.03(-0.13, 0.07) | 0.616 |
|  | cg12881363 | 0.07(-0.06, 0.19) | 0.293 |
|  | cg17657037 | -0.11(-0.21, -0.01) | 0.032 |

Using available follow-up data, we were able to test the temporality between overall school QoL and DNAm (i.e., does wave 3 environment predict follow-up DNAm, and vice versa). We found no significant association in either direction (wave 3 school QoL > follow-up DNAm: -0.0003 [-0.10, 0.10], *p*=0.95; wave 3 DNAm > follow-up school QoL: -4.72 [-9.85, 0.41], *p*=0.07). For household income, there was no follow-up data available to confirm temporal directionality of this significant finding.

*Bold=FDR < 0.05. BMI=Body Mass Index; QoL=Quality of Life*.

The moderating effect of ADHD status on the association between early life and childhood environments and intra-individual change in DNAm was also examined. Comparisons were between Remitted and Persistent ADHD groups to Controls, and between ADHD diagnostic outcome (Remitted ADHD vs. Persistent ADHD).

Across these group comparisons, we identified a significant moderating effect of ADHD status on the association between household income and intra-individual change in cg02208776 DNAm, and this was the only environmental factor that survived FDR correction (**Supplementary Table 6**). Relative to Controls, higher household income in Remitted ADHD was significantly associated with a greater intra-individual decline in cg02208776 DNAm. Comparisons between Persistent ADHD and Controls did not reach statistical significance (*uncorrected p*=0.67). In a separate model comparing Remitted ADHD to the Persistent ADHD group, a significant negative three-way interaction suggests groups had an inverse direction of association.

Stratified LMMs confirmed these direction of these associations within each diagnostic group, though only the result for the Remitted ADHD group reached nominal significance. Patterns of association within each group are shown in **Figure 5**, and present how higher household incomes were associated with an intra-individual decline in cg02208776 DNAm in Remitted ADHD (Estimate [95% CI]=-0.34[-0.54, 0.13], *uncorrected-p*=0.010) and, to a lesser extent, Controls (-0.03[-0.14, 0.08], *uncorrected-p*=0.58). Persistent ADHD had an opposing direction of association, with a higher household income linked to individual increases in DNAm over time (0.04[-0.05, 0.13], *uncorrected- p*=0.42).

**Figure 5.**
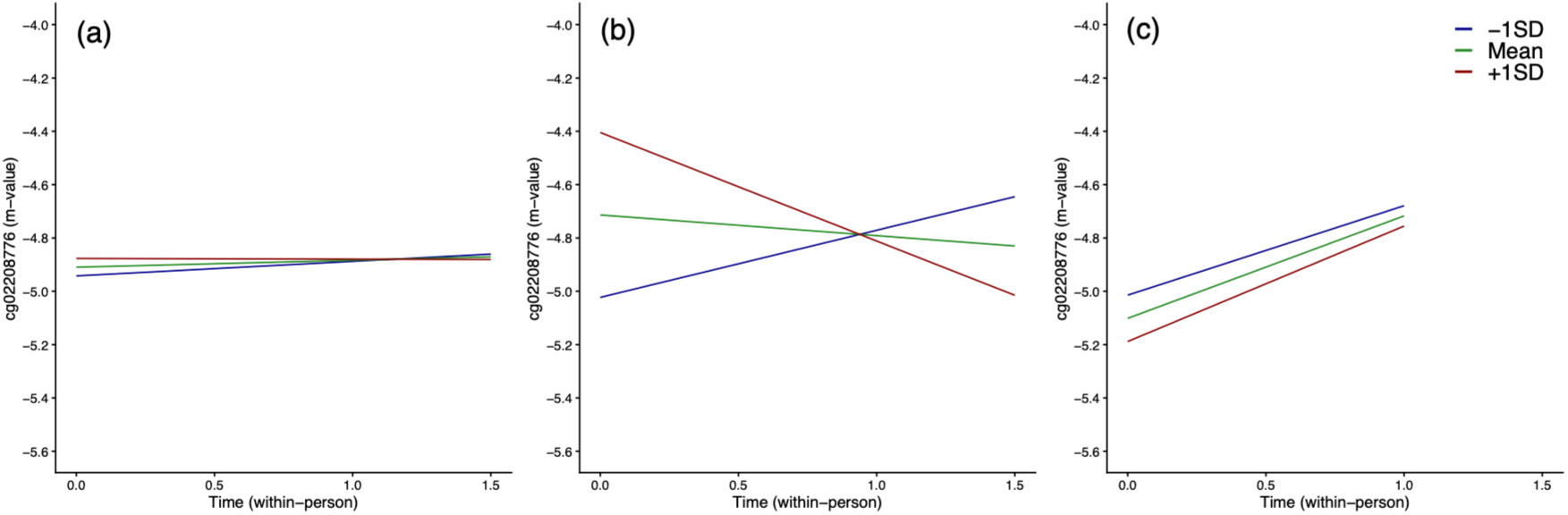
Results from stratified linear mixed models confirming direction of associations between household income and intra-individual change in cg02208776 DNA methylation (DNAm) within each diagnostic group, including (a) Controls, (b) Remitted ADHD and (c) Persistent ADHD. *Legend: Time represents a person-mean centred change, calculated as the difference between each participant’s chronological age at each wave and their age at the first assessment (age at any wave – age at first wave), which was used to model within-person change in DNAm*.

## DISCUSSION

This study extends previous findings (Wrigglesworth et al., 2026) which identified 12 CpGs with varying longitudinal DNAm across ADHD groups, by examining genetic and environmental influences. Our cross-sectional work found 19 single nucleotide polymorphisms (SNPs) that explained a large proportion of variation in DNAm at five CpGs (cg02208776, cg12881363, cg17601216, cg17657037, cg21443143), which were located near (*cis-meQTL*), or further away (*trans-meQTL*). We also identified 17 early life and childhood environmental factors associated with ADHD status and DNAm. We used longitudinal analyses to test whether these environmental factors were also associated with intra-individual change in DNAm, and if these effects varied by ADHD status. Across the whole cohort, a higher school quality of life was associated with an intra-individual decrease in cg11124426 DNAm over time. We also highlight in the whole cohort an association between higher amounts of household income and decreases in intra-individual change in DNAm at cg02208776 (*HSPE1;HSPD1*). This relationship was moderated by ADHD status, extending our previous finding of elevated DNAm in Remitted ADHD relative to Controls (Wrigglesworth et al., 2026). Notably, higher-income individuals with Remitted ADHD showed greater decreases in DNAm over time than Controls, which contrasted patterns observed in Persistent ADHD, thus emphasising income as a protective influence on epigenetic change in Remitted ADHD.

Genetic and environmental variation are well established contributors to ADHD risk, though the underlying biological mechanisms have remained poorly understood. When looking at ADHD diagnostic outcome, this study demonstrated that DNAm at 5 of the 12 CpGs were influenced by genetic variation. This is consistent with previous epigenetic studies in ADHD, which also demonstrate genetic control of differentially DNAm sites, though at CpG’s not investigated here (Goodman et al., 2020; Mooney et al., 2020). We identified 19 independent meQTLs, most of which were trans-chromosomal. Many of these trans associations were polygenic, with groups of variants converging on one of two CpG sites (cg02208776 and cg17657037). By contrast, all cis-meQTLs had a one-to-one relationship with CpG DNAm. This pattern corroborates previous findings (Gaunt et al., 2016), which showed variants closer to a DNAm site tend to have a stronger effect on DNAm than more distal variants; therefore, suggesting trans-meQTLs must act collectively to influence DNAm. However, it should be noted that while we accounted for clumping linkage disequilibrium (LD, i.e., a non-random association between alleles at different loci) on the same chromosome, we did not correct for LD between SNPs found on non-homologous chromosomes (Huang et al., 2023; Lv et al., 2026).

The genetic variants discovered here were mostly linked to protein-coding genes, among which four are important for glycosylation (*ST3GAL1*, *B3GLCT*) or lipid metabolism (*ELOVL6*, *NUDT12)* – two biological processes found to be altered in ADHD (Kianickova et al., 2023; Pivac et al., 2011; Predescu et al., 2024). Genome-wide association studies also link these and other discovered protein-coding genes to brain structures (Alliey-Rodriguez et al., 2019; Fan et al., 2022; N. Liu et al., 2023; van der Meer et al., 2021), and ADHD (Mick et al., 2010). Notably, previous work in this cohort (Dipnall et al., 2024) reported nominal ADHD-DNAm associations at a *ST3GAL1*-annotated CpG, which the current study linked to a cis-meQTL . Lastly, one trans-genetic variant was identified (rs897378832), annotated to a long-non-coding RNA (*LINC01440*), which defines a group of molecules known for their crosstalk with DNAm to regulate gene expression (Yang et al., 2022). Whether these findings represent a genetic modulation of downstream DNAm patterns, or the influence of DNAm on variant transcription would require further investigation.

Whilst the genetic influence on DNAm can be relatively stable across development, the contribution of environment is expected to fluctuate over the life span (Gaunt et al., 2016). We first identified 17 clinically relevant environmental predictors of DNAm, characterising the individual (e.g., child’s health including quality of life, physical health, BMI, psychiatric history and sleep quality) and their surrounding environment (air pollution, perinatal environments, parental mental health history, parenting, stressful life events, income, teacher relationship), which replicate consensus ADHD risk (Faraone et al., 2021). Apart from childhood psychiatric history, all these factors were cross-sectionally associated with DNAm at target sites, emphasising an environmental influence on ADHD that might operate through DNAm. From this selection of environmental predictors, we also identified two FDR-significant longitudinal associations across the whole cohort. First, better school functioning was associated with intra-individual reductions in cg11124426 DNAm, a site where a lower average DNAm level was associated with Remitted ADHD, when compared to Controls in our previous study (Wrigglesworth et al., 2026). Secondly, higher household income was also associated with reductions in cg02208776 DNAm over time, for which a higher average DNAm level at this site was also associated with Remitted ADHD, when compared to Controls in our previous study (Wrigglesworth et al., 2026). School functioning, representing indices of quality of school engagement (e.g., parent reports of how much a child pays attention in class or keeps up with schoolwork), has been associated with mental health problems, including ADHD (Lee et al., 2016; Wen et al., 2026). Interestingly we identified no significant moderating effect of ADHD status on this relationship, but rather represented a link across all groups. While there are currently no studies to our knowledge that have investigated a relationship between school functioning and DNAm, there are a few candidate and epigenome wide studies (EWAS) on related areas that tend to overlap with school functioning, including attention and academic performance (Burenkova et al., 2024; Camerota et al., 2024; Ehlinger et al., 2023). Evidence from these studies corroborate our negative association through links between early life DNAm (infant) and early childhood attentional problems (at 2 years of age) (Camerota et al., 2024), and a cross-sectional relationship between DNAm and measures of academic performance in middle-childhood (writing composite scores, picture vocabulary) (Burenkova et al., 2024). However, positive relationships with attention and academic performance have also been reported (Ehlinger et al., 2023), even within these aforementioned studies. Given the scarcity of epigenetic research on school functioning, there remains a need for longitudinal investigations to validate our finding.

A higher household income was linked to an intra-individual decrease in cg02208776 DNAm (*HSPE1;HSPD1*) over time, and these associations varied between Remitted ADHD and Controls, and by clinical outcome (Remitted ADHD vs. Persistent ADHD). While there have been several studies which associate composite measures of socioeconomic (SES) status with DNAm (Beach et al., 2014; Beach et al., 2016; Laubach et al., 2019; Swartz et al., 2017), few have investigated individual household income. Among three known studies on household income, only one prospective EWAS identified CpGs where early childhood income-per-dependent was associated with later childhood DNAm, representing CpGs with either a negative or positive association with income (Bush et al., 2018). The two other studies, one candidate and one EWAS, reported null findings (Coker et al., 2018; J. Liu et al., 2023).

We also identified significant ADHD group differences in the relationship with household income, highlighting a greater reduction in intra-individual change in cg02208776 DNAm in Remitted ADHD at higher levels of income, compared to Persistent ADHD and Controls. To our knowledge this is the first study to test these comparisons, though there is evidence associating higher incomes with a reduced risk of ADHD, and a better clinical outcome, which support our study finding (Kim et al., 2019; Law et al., 2014). Moreover, neuroimaging studies demonstrate how income can influence brain structures (Machlin et al., 2020; Pan et al., 2026; Xie et al., 2026), with links between income and ADHD related to the prefrontal working memory system (Carlsson et al., 2021; Hackman et al., 2010). What cannot be inferred from the current study findings is whether household income is directly associated with intra-individual change in DNAm, or if there is a third mediating factor through which income can influence epigenetic change. One potential explanation may be that individuals with higher incomes are more financially capable to invest in ADHD medication, better nutrition, housing, schools and recreation (Adler & Newman, 2002). Moreover, our finding also shows that genetics explains variance at this CpG site. Thus, we cannot rule out the possibility of familial confounding, whereby associations are driven by genetic differences across the population. Statistical adjustment for genetic variation would have addressed this potential confounding, though this may not have been feasible given our small sample size. From a recent registered systematic review, findings highlight evidence showing a link between income and ADHD, even after adjusting for genetic differences (Carlsson et al., 2021).

This study has several strengths. First and foremost, the current study is one of the first to investigate environmental associations with longitudinal DNAm, and whether these vary by clinical outcomes of remittance and persistence in ADHD. We used prior knowledge from an international consensus statement (Faraone et al., 2021) to select environmental correlates of ADHD risk. In addition to investigating known environmental predictors of ADHD, we addressed its genetic contribution (Demontis et al., 2019) by testing whether genetic variants were associated with ADHD-related DNAm – which we assessed through our meQTL analysis.

Notwithstanding these strengths are several limitations that warrant consideration. While this cohort had epigenetic data across three waves, most individuals attended two assessments, and across a limited age range. This is also applicable to historical exposures of early life and childhood environment which were collected from questionnaires conducted at two waves prior and at the same time of biological sampling, though not all participants had attended all three visits. Moreover, 40% of the whole environmental cohort did not contribute longitudinal epigenetic data. Although these individuals were comparable to the remaining cohort in baseline characteristics, they reported significantly lower QoL scores, including school functioning; more peer relational victimisation, and were more likely to have a history of internalising disorder. Diagnostic status was assessed at recruitment, and wave 3 and 5, but was not reassessed at the intervening waves. Consequently, while participants with persistent ADHD or without ADHD continued to meet their respective diagnostic criteria at the final wave, it was not possible to determine a more precise timing of remission within a 36month window. Another limitation is that we did not adjust environmental analyses for potential genetic confounding. This is particularly relevant for home environment variables like household income. However, given our relatively small sample size, this additional adjustment may not have been feasible. Lastly, these findings are limited to epigenetic markers measured from saliva, and therefore cannot inform directly on DNAm in brain tissue. Thus, while some studies report relative stability of DNAm across tissue types (Nishitani et al., 2023), patterns are highly tissue- and cell-specific, and these findings should be interpreted with caution.

In conclusion, in children and adolescents with and without ADHD we extend the current literature with novel findings that highlight genetic control over some, though not all epigenetic markers of ADHD. Moreover, we provide evidence which suggests better school functioning and higher levels of household income are associated with decreases in DNA methylation over time, with this negative association most relevant to individuals with remitted ADHD symptoms. Given the novelty of these findings, which were conducted on a relatively small cohort of individuals assessed over a short duration, it is necessary to validate these findings using longitudinal studies with larger cohorts investigated over longer periods of time. In addition, future studies would benefit in considering the interplay of genes on physical and social environments.

## DATA AVAILABILITY STATEMENT

Data from the Children’s Attention Project cohort are available via Lifecourse: https://lifecourse.melbournechildrens.com/cohorts/cap-and-nicap/. The data that support the findings of this study are available on request from the corresponding author, upon reasonable request.

## Supporting information

Supplementary methods and results

## ACKNOWLEDGEMENTS

A special thanks to all the children and families for their participation in this study. We also thank Professor Jeffrey M. Craig for his role as co-investigator on NICAP and for his assistance with the original EWAS work. Professor Craig was unfortunately unable to contribute to the current study. This work was supported by the MASSIVE HPC facility (www.massive.org.au).

## AUTHOR CONTRIBUTION

J.W.: data curation, conceptualization, methodology, formal analysis, writing – original draft, and visualization. V.C.: methodology, formal analysis, writing – original draft, writing – review & editing. P.D.F.: data curation, methodology, writing – review and editing. V.A.: writing – review & editing. J.M.C.: project administration, resources, writing – review & editing. T.J.S.: supervision, project administration, resources, conceptualization, methodology, writing – review & editing.

