## Supplementary methods and results for "Genetic and environmental factors associated with DNA methylation across childhood and adolescence in individuals with and without attention-deficit/hyperactivity disorder"

#### Environmental exposures

*Variable coding*

Environmental data collected across Waves 1-3, were coded to represent exposures occurring before or concurrent with Wave 3 DNAm, either using an '*Absolute Exposure*' or '*Overall Exposure*' level approach. Absolute Exposure was defined as a positive code if the condition was measured at *any* timepoint (Wave 1, 2, or 3), and this coding was applied to perinatal factors, parental mental health history, child medication history, and all stressful life event exposures (**Supplementary Table 1**). For all other environmental variables, we use the Overall Exposure to represent the median level of exposure measured across the first three waves.

*Variable quality control*

Environmental variables underwent quality control (QC) procedures after coding. This includes control for missing data, frequency imbalance (for categorical variables), skewness (for continuous variables), near-zero variances and multicollinearity. Variables were removed from the analyses if they fell into one of the following exclusionary criteria: 1) more than 10% of missing data; 2) frequency leading to less than 10% of observation in the minority group; 3) strong skewness defined as <-1.6 or >1.6; iv) near-zero variances (defined as less than 10% of distinct values); 4) multicollinearity (defined as Pearson r> 0.90).

#### Genetic variants

DNA extracted from saliva were genotyped using the Illumina Global Screening Array (GSA-MD-24v1-0; Illumina, San Diego, CA, USA) and initially pre-processed at the Stanley Center for Psychiatric Disease, Broad Institute (Cambridge, MA, USA). Initial QC procedures were executed using the Ricopili pipeline (Lam et al., 2020), and involved three consecutive QC runs at both the sample and variant levels.

Sample-level filtering commenced after the removal of single nucleotide polymorphisms (SNPs) with a call rate < 95%. Individuals with a call rate < 0.98 or inbreeding coefficient (F) > 0.20

were initially excluded. Biological sex was verified using X-chromosome data, and samples with discordant sex information (defined as male $F < 0.5$ or female $F > 0.5$) were removed.

At the variant-level, SNPs were excluded if they met any of the following criteria: 1) call rate < 0.98; 2) missingness difference between cases and controls > 0.02; 3) monomorphic (indicating low variance), or 4) deviated from the Hardy-Weinberg Equilibrium (HWE) at thresholds of $p < 1*10^{-6}$ (Controls) or $p < 1*10^{-10}$ (Cases). The performance of each QC iteration was evaluated using an uncorrected GWAS genomic inflation value. To mitigate confounding by relatedness, pairs demonstrating a pi-hat > 0.2 were identified, and one individual from each pair was excluded, with a priority placed on retaining cases.

Population stratification was addressed using a Principal Component Analysis (PCA), projecting the cohort onto the 1000 Genomes Project reference panel (1000 Genomes Project Consortium et al., 2015). Samples clustering within the European ancestry group were retained for downstream analysis.

Missing data underwent imputation before the final automated QC procedures were performed. Here, the *CheckVCF* tool was first used to perform an initial allele check on our QC GRCh37 (hg19) build dataset, and corrected allele swaps using the *bcftools* software (Danecek et al., 2021). The verified dataset was subsequently submitted to the TOPMed Imputation Server (https://imputation.biodatacatalyst.nhlbi.nih.gov) for phasing and imputation, where it was automatically lifted over to the GRCh38 (hg38) genome build. Haplotype phasing was conducted using Eagle v2.4 (Loh et al., 2016), and genotype imputation was performed using Minimac4 (https://github.com/statgen/Minimac4) against the NHLBI TOPMed Reference Panel (Version r3, GRCh38). Automated post-imputation QC was performed by the server, which retained variants with an imputation quality $R^2 \geq 0.3$ and a minor allele frequency (MAF) $\geq 0.05$. A total of 8,231,947 variants were retained for downstream analyses, with the input data demonstrating a high concordance with the reference panel, reflecting a 96.02% reference overlap.

**Statistical analyses**

1) *Methylation quantitative trait loci (meQTL) analyses*

*Custom Linkage Disequilibrium Clumping algorithm*

To distinguish independent genetic signals from associations that can be attributed to the same causal variant on a homologous chromosome through linkage disequilibrium (LD), we applied a custom greedy LD-clumping algorithm to each CpG site with an FDR-significant association. The custom algorithm follows similar logic to PLINK (Purcell et al., 2007), where all FDR-significant associations, for any given probe, were first ranked by p-value from the most to least significant. The most significant SNP was designated the 'lead' or index SNP for a new clump. Pairwise squared Pearson correlation coefficient ($r^2$) values were then computed (directly from the imputed dosage matrix of study participants, $n \approx 113$) between the lead SNP and all remaining significant SNPs located on the same chromosome within a ±500 kb physical window. Any SNP with $r^2 \geq 0.10$ relative to the lead SNP was assigned to that clump and removed from subsequent consideration. This step was iterated until no further per-CpG-significant FDR-associations remained for that site. The $r^2$ threshold of 0.10 and the 500 kb window are consistent with standard practice in genome-wide association studies and represent a balance between excluding correlated signals and retaining genuinely independent loci.

Each surviving lead SNP represents a distinct independent genetic signal for the corresponding CpG. For each lead SNP, the magnitude of genetic effect on CpG DNA methylation (DNAm) is characterized by the regression beta coefficient (estimated change in DNAm M-value per additional copy of the effect allele under the additive model) and by the partial $R^2$ (estimated increment in explained variance when SNP term was added to a covariate-only, ordinary least-squares model). This partial $R^2$ quantifies the unique proportion of DNAm variance attributable to the SNP genotype after accounting for age, sex, ADHD status, batch, epithelial cell-proportion, and population stratification.

2) <u>*Environmental predictors of DNA methylation*</u>

Cross-sectional analyses used a two-step approach to identify clinically relevant environmental predictors of Wave 3 DNAm. This approach sequentially applied Least Absolute Shrinkage and Selection Operator (LASSO) regression models: the first step identified nominally significant environmental predictors of ADHD status, and the second step determined whether these environmental factors are also associated with DNAm. Each step is detailed in the subsections below.

*Step 1. Environmental predictors of ADHD status*

The first step reduced the large number of QC surviving environmental factors (**Supplementary Table 1**) into a set of predictors of ADHD status. Here we ran three separate LASSO regression models, each predicting a pairwise comparison: 1) Remitted ADHD vs. Controls; 2) Persistent ADHD vs. Controls; and 3) Remitted ADHD vs. Persistent ADHD.

Models were implemented using the *glmnet* package in R (Friedman et al., 2010), and all continuous predictors were standardised prior to modelling. For each LASSO procedure, we applied 10-fold cross-validation to identify the optimal penalty parameter, which minimised prediction error. We pre-specified child age and sex as necessary covariates for all models, and these were kept unpenalized. All remaining environmental predictors were subject to the regularisation penalty (penalty factor=1). For interpretation purposes, the environmental predictors with a non-zero coefficients (at lambda_min) were carried forward into an unregularized binary logistic regression model (*glm*) to extract associated Odds Ratios (ORs).

*Step 2. Environmental predictors of wave 3 DNA methylation*

After identifying significant environmental factors of ADHD status in Step 1, we examined which of these predictors were also associated with Wave 3 DNAm, at any of the 12 CpG sites. A similar LASSO procedure was run separately for each CpG, and used their residualised DNAm values as the outcome to account for technical batch (i.e., plate), epithelial cell-proportion, child age and sex. Residualised DNAm for each CpG site was derived using the linear model fitting function and residual extraction from the *limma* package (Phipson et al., 2016). LASSO models were fitted using a Gaussian family distribution with 10-fold cross-validation to optimally select predictors associated with residualised DNAm levels at each CpG site.

### SUPPLEMENTARY RESULTS

#### Environmental predictors of DNA methylation

<u>*Cross-sectional DNA methylation in late childhood*</u>

We first examined whether there are clinically relevant early life and childhood environmental predictors of Wave 3 DNAm. Across two steps of analysis, we applied LASSO regression models to identify environmental predictors associated with ADHD status (Step 1), and that are also associated with DNAm (Step 2). We detail these results in the sections below.

*Step 1. Environmental predictors of ADHD status*

A total of 23 environmental factors were selected as predictors of ADHD status: 14 predicted Remitted ADHD vs. Controls; 10 predicted Persistent ADHD vs. Controls, and 6 predicted Remitted ADHD to Persistent ADHD (**Supplementary Table 3**). These predictors covered a range of environmental categories, including *Pollution* (n=1), *Home environment* (n=4), *School environment* (n=1), *Perinatal* (n=1), and *Stressful life events* (n=4), with most predictors relating to the *Child's health* (n=12).

From pairwise comparisons between Remitted ADHD and Controls, there were increased odds of Remitted ADHD associated with a higher median household income (OR=2.21); history of psychotropic medication use (OR=2.16); a higher overall-reported health status score (OR=1.93); history of sleep initiation (OR=1.85) and maintenance (OR=1.35) problems; a history of externalised disorders (OR=1.78); the experience of moving house (OR=1.77); a higher median Body Mass Index (BMI) (OR=1.64) and a higher median level of PM2.5 exposure (OR=1.49). Conversely, a decreased odds of Remitted ADHD was associated with a higher median child quality of life (QoL) scores related to school functioning (OR=0.18); parental history of any psychiatric disorders (OR=0.60); a stressful life event of experiencing a death of a close friend/family member (OR=0.68); a higher median Teacher-Student Relationship total score (OR=0.89), and a history of daytime sleepiness (OR=0.99).

When comparing Persistent ADHD to Controls, an increased odds of Persistent ADHD was linked to the presence of any sleep problem (OR=2.33); higher median levels of angry parenting (OR=2.24); a stressful life event of experiencing death of partner or child's grandparent (OR=1.47); a history of externalised (OR=1.27) and/or internalised disorders (OR=1.26); history of anxiety during pregnancy (OR=1.24), and the use of any psychotropic medication (OR=1.15). The odds of Persistent ADHD were reduced at higher child QoL scores related to school functioning (OR=0.24) and psychosocial wellbeing (OR=0.742), and higher median levels of the student-teacher relationship total score (OR=0.62).

Lastly, LASSO was used to identify early life and childhood environmental predictors of Persistent ADHD, using a pairwise comparison with the Remitted ADHD group. Here, increased odds of Persistent ADHD were associated with higher median levels of parental ADHD symptoms severity (OR=1.35) and angry parenting (OR=1.18). The odds of Persistent ADHD were reduced at higher median QoL scores related to emotional (OR=0.56) and school functioning (OR=0.77) and a higher BMI (OR=0.62).

*Step 2. Environmental predictors of Wave 3 DNA methylation*

The above-mentioned environmental predictors of ADHD status (Step 1) were then analysed as correlates of Wave 3 DNAm at each 12 CpG site, using a similar LASSO procedure. Here we identified environmental predictors associated with DNAm at 6 CpGs (cg02208776 [*HSPE1;HSPD1*], cg02968116, cg11124426, cg12577151 [*ACAD9*], cg1281363 [*MOV10L1*], and cg17657037 [*ITGB1BP3*]), which we detail in **Supplementary Table 4**. These features covered environmental categories of *Child's health* (n=7); *Home environment* (n=4); *Perinatal* (n=1); *Pollution* (PM2.5); *School environment* (n=1) and *Stressful life events* (n=3). Both cg1124426 and cg17657037 (*ITGB1BP3*) were linked to the most environmental predictors, while cg02208776 and cg12577151 (*ACAD9*) were associated with the fewest (1 feature each).

For cg1124426, we identified nine predictors with a positive association: median levels of school functioning; PM2.5 exposure; angry parenting; student-teacher total relationship score, and the experience of a close family member being ill/injured or assaulted (**Supplementary Table 4**). Conversely, negative associations for cg11124426 included a history of anxiety during pregnancy, experiencing death of a close friend/family member, a history of problems maintaining sleep, and median levels of QoL scores related to emotional functioning.

A total of 13 environmental predictors were associated with DNAm at cg17657037. Here, 6 predictors were negatively associated with DNAm, including median levels of QoL scores related to psychosocial wellbeing; parent-rated child's overall health score; experiencing problems initiating sleep, or stressful life events related to moving house; median levels of household income, and angry parenting. Conversely, the remaining 7 predictors were positively associated with DNAm of cg17657037, including higher median levels of QoL scores related to emotional functioning; BMI; parent ADHD symptom severity, or parents history of any mental health disorder; median levels of student-teacher total relationship, and experiencing illness/injury or assault of a close family member or death of a close friend/family member (**Supplementary Table 4**).

CpG sites with the fewest number of predictors were cg02208776 and cg12577151, each with a single environmental feature linked to DNAm. Here, higher cg02208776 DNAm was associated with a higher median household income. For cg12577151, higher median levels of PM2.5 were associated with a higher DNAm (**Supplementary Table 4**).

*Longitudinal change in DNA methylation*

Fifty-one participants were not eligible for this portion of study as they had only provided a single biological sample. These individuals were otherwise similar across most demographics aside from having a lower overall QoL score for physical functioning (median 80.0 vs. 90.6; Kruskal-Wallis (KW) $p = 0.004$), school functioning (median 55.0 vs. 70.0; KW $p = 0.019$), psychosocial health summary score (median 65.0 vs. 76.7; KW $p = 0.024$), and total scale score (median 71.7 vs. 80.0, KW $p = 0.004$); a higher overall score for experiencing peer relational victimisation (median 2.0 vs. 1.7, KW $p = 0.029$), and a greater number of participants with a history of internalising disorder (31.4% vs. 14.7%, Fisher exact $p = 0.029$).

**SUPPLEMENTARY TABLES**

**Supplementary Table 1.** Listing of assessed environmental exposures.

| Study category | Exposure | Data type |
|---|---|---|
| *Child's health* | QoL: physical functioning | Overall Exposure |
| | QoL: emotional functioning | Overall Exposure |
| | QoL: social functioning | Overall Exposure |
| | QoL: school functioning | Overall Exposure |
| | QoL: psychosocial health summary school | Overall Exposure |
| | QoL: total scale score | Overall Exposure |
| | Body Mass Index (BMI) | Overall Exposure |
| | Parents rating of child's overall health | Overall Exposure |
| | History of internalising disorder | Absolute Exposure |
| | History of externalising disorder | Absolute Exposure |
| | Taking ADHD stimulant medications | Absolute Exposure |
| | Taking any psychotropic medication | Absolute Exposure |
| | Respiratory problems during sleep | Absolute Exposure |
| | Problems initiating sleep | Absolute Exposure |
| | Problems maintaining sleep | Absolute Exposure |
| | Daytime sleepiness | Absolute Exposure |
| | Any sleep problems | Absolute Exposure |
| *Home environment* | Parental relative poverty | Overall Exposure |
| | Parent psychosocial well-being | Overall Exposure |
| | Parent ADHD symptom severity | Overall Exposure |
| | Parent positive affect | Overall Exposure |
| | Household income | Overall Exposure |
| | Consistent parenting | Overall Exposure |
| | Angry parenting | Overall Exposure |
| | Parental warmth | Overall Exposure |
| | Parents history of anxiety disorders | Absolute Exposure |
| | Parents history of mood disorders | Absolute Exposure |
| | Parents history of any mental health disorders | Absolute Exposure |
| *Perinatal* | Intensive care at birth | Absolute Exposure |
| | Maternal diabetes during pregnancy | Absolute Exposure |
| | Maternal high blood pressure during pregnancy | Absolute Exposure |
| | Maternal anxiety during pregnancy | Absolute Exposure |
| | Maternal stressful life event during pregnancy | Absolute Exposure |
| | Maternal post-natal depression | Absolute Exposure |
| | Maternal alcohol during pregnancy | Absolute Exposure |
| | Maternal smoking during pregnancy | Absolute Exposure |
| | Low-birth weight (<2500g) | Absolute Exposure |
| | Premature birth (<37weeks) | Absolute Exposure |
| *Pollution* | Nitrogen Dioxide (NO2) pollution | Overall Exposure |
| | Air particulate matter (pm) 2.5 | Overall Exposure |
| *School environment* | SEQ: Peer relational victimisation | Overall Exposure |
| | SEQ: Peer physical victimisation | Overall Exposure |
| | Student teacher relationship - closeness | Overall Exposure |
| | Student teacher relationship - total | Overall Exposure |
| | Experience of bullying | Absolute Exposure |
| *Stressful life events* | Parent illness/injury/assault | Absolute Exposure |
| | Close family member illness/injury/assault | Absolute Exposure |

| | | |
|---|---|---|
| | Death of partner or child's grandparent | Absolute Exposure |
| | Death of close friend/relative | Absolute Exposure |
| | Parent separation/relationship end | Absolute Exposure |
| | Serious problem with a relative/close friend/neighbour | Absolute Exposure |
| | Parent/partner lost job | Absolute Exposure |
| | Major financial difficulties | Absolute Exposure |
| | Parent/close family member had trouble with police/ court appearance | Absolute Exposure |
| | Valuables lost/stolen | Absolute Exposure |
| | Moved house | Absolute Exposure |
| | Parent becoming pregnant/having a child | Absolute Exposure |
| | Any stressful event | Absolute Exposure |

*'Overall Exposure'=the median level of exposure measured across the first three assessment waves; 'Absolute Exposure'=binary categorical variables with a positive code (1, 'Yes') if the condition was measured at* any *time point. SEQ=Social Experience Questionnaires; QoL=Quality of Life.*

**Supplementary Table 2.** Summary of early life and childhood environmental predictors of ADHD status and wave 3 DNAm used in the two-step LASSO regression approach.

| Environmental exposures | Control (N=62) | Remitted (N=26) | Persistent (N=38) | Total (N=126) |
|---|---|---|---|---|
| Chronological age, mean(SD) | 10.4(0.4) | 10.4(0.4) | 10.4(0.5) | 10.4(0.5) |
| Female, Yes(%) | 26(42%) | 11(42%) | 9(24%) | 46(37%) |
| **Child's health:** | | | | |
| QOL: physical functioning, mean(SD) | 90.0(10.1) | 74.7(19.2) | 82.1(14.7) | 84.5(15.0) |
| QOL: emotional functioning, mean(SD) | 76.6(13.9) | 62.2(20.3) | 56.2(15.2) | 67.5(18.2) |
| QOL: social functioning, mean(SD) | 87.7(12.7) | 67.5(20.9) | 63.0(19.3) | 76.1(20.3) |
| QOL: school functioning, mean(SD) | 80.8(15.5) | 56.2(10.7) | 46.9(16.5) | 65.5(21.5) |
| QOL: psychosocial health summary score, mean(SD) | 80.9(10.9) | 60.8(13.9) | 56.6(13.0) | 69.4(16.7) |
| QOL: total scale score, mean(SD) | 84.0(9.4) | 65.0(13.2) | 64.3(13.5) | 74.1(15.0) |
| BMI, mean(SD) | 17.5(2.4) | 19.7(3.7) | 17.9(3.3) | 18.1(3.1) |
| Parents rating of child's overall health, mean(SD) | 4.5(0.7) | 4.2(0.5) | 4.5(0.6) | 4.4(0.7) |
| History of internalising disorder, Yes(%) | 6(10%) | 5(19%) | 16(42%) | 27(21%) |
| History of externalising disorder, Yes(%) | 6(10%) | 16(62%) | 24(63%) | 46(37%) |
| Taking ADHD stimulant medications, Yes(%) | 0(0%) | 4(15%) | 11(29%) | 15(12%) |
| Taking any psychotropic medications, Yes(%) | 0(0%) | 6(23%) | 13(34%) | 19(15%) |
| Respiratory problems during sleep, Yes(%) | 11(18%) | 6(23%) | 8(21%) | 25(20%) |
| Problems initiating sleep, Yes(%) | 24(39%) | 20(77%) | 31(82%) | 75(60%) |
| Problems maintaining sleep, Yes(%) | 14(23%) | 16(62%) | 20(53%) | 50(40%) |
| Daytime sleepiness, Yes(%) | 18(29%) | 18(69%) | 23 (60.5%) | 59 (46.8%) |
| Any sleep problems, Yes(%) | 34(55%) | 24(92%) | 36 (94.7%) | 94 (74.6%) |
| **Home environment:** | | | | |
| Parental relative poverty, mean(SD) | 1024.5(46.2) | 1017.7(38.4) | 1017.2(41.0) | 1020.9(43.0) |
| Parent psychosocial well-being, Yes(%) | 2.5(2.6) | 4.0(3.6) | 5.6(4.7) | 3.7(3.8) |
| Parent ADHD symptom severity, mean(SD) | 7.9(4.2) | 10.3(5.0) | 12.2(6.0) | 9.7(5.3) |
| Parent positive affect, mean(SD) | 9.5(2.4) | 7.4(2.3) | 8.4(2.6) | 8.7(2.6) |
| Household income (AUD), mean(SD) | 84193.5(20085.8) | 84230.8(20478.9) | 74736.8(24849.5) | 81349.2(21967.4) |
| Consistent parenting, mean(SD) | 25.2(3.6) | 24.3(2.8) | 24.0(3.7) | 24.7(3.5) |
| Angry parenting, mean(SD) | 9.7(2.3) | 12.1(3.2) | 13.0(3.5) | 11.2(3.2) |
| Parental warmth, mean(SD) | 26.0(2.9) | 24.7(4.5) | 25.2(3.8) | 25.5(3.6) |
| Parent history of anxiety disorders, Yes(%) | 12(19%) | 6(23%) | 9(24%) | 27(21%) |

| | | | | |
|---|---|---|---|---|
| Parent history of mood disorders, Yes(%) | 19(31%) | 13(50%) | 17(45%) | 49(39%) |
| Parent history of any mental health disorders, Yes(%) | 23(37%) | 13(50%) | 17(45%) | 53(42%) |
| **Perinatal:** | | | | |
| Intensive care at birth, Yes(%) | 11(18%) | 6(23%) | 8(21%) | 25(20%) |
| Maternal diabetes during pregnancy, Yes(%) | 5(8%) | 1(4%) | 4(11%) | 10(8%) |
| Maternal HBP during pregnancy, Yes(%) | 4(7%) | 4(15%) | 1(3%) | 9(7%) |
| Maternal anxiety during pregnancy, Yes(%) | 10(16%) | 6(23%) | 13(34%) | 29(23%) |
| Maternal SLE during pregnancy, Yes(%) | 14(23%) | 6(23%) | 14(37%) | 34(27%) |
| Maternal post-natal depression, Yes(%) | 9(15%) | 5(19%) | 8(21%) | 22(18%) |
| Maternal smoking during pregnancy, Yes(%) | 10(16%) | 5(19%) | 8(21%) | 23(18%) |
| Maternal alcohol during pregnancy, Yes(%) | 21(34%) | 9(35%) | 13(34%) | 43(34%) |
| Low-birth weight (<2500g), Yes(%) | 1(2%) | 3(12%) | 4(11%) | 8(6%) |
| Premature birth (<37wks), Yes(%) | 4(7%) | 5(19%) | 5(13%) | 14(11%) |
| **Pollution:** | | | | |
| Nitrogen Dioxide (NO2) pollution, mean(SD) | 1.5(0.4) | 1.4(0.4) | 1.5(0.3) | 1.5(0.4) |
| Air particulate matter (pm) 2.5, mean(SD) | 5.5(0.7) | 5.5(0.7) | 5.4(0.6) | 5.5(0.7) |
| **School environment:** | | | | |
| SEQ: Peer relational victimisation, mean(SD) | 1.5(0.5) | 2.2(10.0) | 2.4(0.9) | 1.9(0.8) |
| SEQ: Peer physical victimisation, mean(SD) | 1.3(0.5) | 1.8(0.8) | 1.9(0.7) | 1.6(0.7) |
| Student-teacher relationship - closeness, mean(SD) | 34.2(3.5) | 32.1(4.2) | 32.5(3.6) | 33.3(3.8) |
| Student-teacher relationship - total, mean(SD) | 63.0(4.4) | 55.9(9.1) | 55.5(6.9) | 59.3(7.3) |
| Experience of bullying, Yes(%) | 30(48%) | 14(54%) | 27(71%) | 71(56%) |
| **Stressful life events:** | | | | |
| Parent illness/injury/assault, Yes(%) | 6(10%) | 4(15%) | 9(24%) | 19(15%) |
| Relative illness/injury/assault, Yes(%) | 25(40%) | 8(31%) | 16(42%) | 49(39%) |
| Death of partner/child's grandparent, Yes(%) | 2(3%) | 3(12%) | 4(11%) | 9(7%) |
| Death of close friend/relative, Yes(%) | 26(42%) | 8(31%) | 18(47%) | 52(41%) |
| Parent separation/relationship end, Yes(%) | 5(8%) | 3(12%) | 6(16%) | 14(11%) |
| Serious problem with family member/close friend/neighbour, Yes(%) | 20(32%) | 11(42%) | 20(53%) | 51(41%) |
| Parent/partner lost job, Yes(%) | 10(16%) | 7(27%) | 6(16%) | 23(18%) |
| Major financial difficulties, Yes(%) | 9(15%) | 6(23%) | 13(34%) | 28(22%) |
| Parent/close relative had trouble police/court appearance, Yes(%) | 6(10%) | 2(8%) | 7(18%) | 15(12%) |
| Valuables lost/stolen, Yes(%) | 4(7%) | 5(19%) | 8(21%) | 17(14%) |

| | | | | |
|---|---|---|---|---|
| Moved house, Yes(%) | 20(32%) | 10(39%) | 14(37%) | 44(35%) |
| Parent becoming pregnant/having a child, Yes(%) | 6(10.0%) | 4(15%) | 9(24%) | 19(15%) |
| Any stressful event, Yes(%) | 39(63%) | 15(58%) | 22(58%) | 76(60%) |

*BMI=Body Mass Index; SEQ=Social Experience Questionnaires; QoL=Quality of Life. AUD=Australian currency.*

**Supplementary Table 3:** Environmental features selected from the first LASSO regression models predicting one of three pairwise ADHD group comparisons.

| Environmental exposure | Remitted ADHD vs. Controls Estimate | Persistent ADHD vs. Controls Estimate | Persistent ADHD vs. Remitted ADHD Estimate |
|---|---|---|---|
| **Child health:** | | | |
| QoL: emotional functioning | | | -0.254 |
| QoL: school functioning | -0.944 | -0.741 | -0.053 |
| QoL: psychosocial health summary school | | -0.477 | |
| BMI | 0.199 | | -0.048 |
| Parents rating of child's overall health | 0.082 | | |
| History of internalising disorder | | 0.063 | |
| History of externalising disorder | 0.288 | 0.157 | |
| Taking any psychotropic medication | 0.306 | 0.040 | |
| Problems initiating sleep | 0.199 | | |
| Problems maintaining sleep | 0.146 | | |
| Daytime sleepiness | 0.066 | | |
| Any sleep problems | | 0.436 | |
| **Home environment:** | | | |
| Parent ADHD symptom severity | | | 0.052 |
| Household income | 0.262 | | |
| Angry parenting | | 0.368 | 0.080 |
| Parents history of any mental health disorders | -0.045 | | |
| **Perinatal:** | | | |
| Maternal anxiety during pregnancy | | 0.027 | |
| **Pollution:** | | | |
| Air particulate matter 2.5 | 0.092 | | |
| **School environment:** | | | |
| Student teacher total relationship | -0.053 | -0.196 | |
| **Stressful life events:** | | | |
| Close family member illness/injury/assault | | | 0.067 |
| Death of partner or child's grandparent | | 0.014 | |
| Death of close friend/family member | -0.049 | | |

| | |
|---|---|
| Moved house | 0.028 |

*Each LASSO model was adjusted for child gender and child age. BMI=Body Mass Index; QoL=Quality of Life.*

**Supplementary Table 4:** Early life and childhood environmental factors selected for the second LASSO regression models predicting wave 3 DNAm at one of the 12 CpG sites.

| Environmental factors | cg02208776 Estimate | cg02968116 Estimate | cg11124426 Estimate | cg12577151 Estimate | cg12881363 Estimate | cg17657037 Estimate |
|---|---|---|---|---|---|---|
| **Child health:** | | | | | | |
| QoL: emotional functioning | | | -0.146 | | | 0.192 |
| QoL: school functioning | | -0.011 | 0.147 | | | -0.018 |
| QoL: psychosocial health summary school | | -0.028 | | | | -0.155 |
| BMI | | | | | -0.060 | 0.065 |
| Parents rating of child's overall health | | | | | | -0.010 |
| Problems initiating sleep | | | | | | -0.079 |
| Problems maintaining sleep | | | -0.060 | | | |
| **Home environment:** | | | | | | |
| Parent ADHD symptom severity | | | | | | 0.021 |
| Household income | 0.042 | | | | | -0.055 |
| Angry parenting | | | 0.0005 | | -0.083 | -0.058 |
| Parents history of any mental health disorders | | | | | | 0.091 |
| **Perinatal:** | | | | | | |
| Maternal anxiety during pregnancy | | | -0.197 | | | |
| **Pollution:** | | | | | | |
| Air particulate matter 2.5 | | | 0.018 | 0.028 | | |
| **School environment:** | | | | | | |
| Student teacher relationship: total | | | 0.027 | | -0.104 | 0.037 |
| **Stressful life events:** | | | | | | |
| Close family member illness/injury/assault | | | 0.044 | | | 0.047 |
| Death of close friend/family member | | | -0.070 | | | 0.018 |
| Moved house | | | | | | -0.082 |

*Each LASSO model was adjusted for child gender and child age. BMI=Body Mass Index; QoL=Quality of Life.*

**Supplementary Table 5.** Clinically relevant early life and childhood environments associated with DNA methylation in late childhood, and further investigated as baseline predictors of longitudinal DNA methylation in participants with two or three waves of epigenetic data (n=75).

| Early life and childhood environments | Control (n=41) | Remitted ADHD (n=15) | Persistent ADHD (n=19) |
|---|---|---|---|
| Chronological age, mean(range) | 10.3(9.5-11.9) | 10.4(9.9-11.0) | 10.3(9.8-11.5) |
| Female, Yes(%) | 12(29%) | 5 (33%) | 6(32%) |
| **Child health:** | | | |
| QOL: physical functioning, mean(SD) | 91.2(10.2) | 77.6(15.6) | 86.4(15.8) |
| QOL: emotional functioning, mean(SD) | 78.2(13.4) | 63.5(17.8) | 58.8(16.3) |
| QOL: psychosocial health, mean(SD) | 81.7(10.1) | 59.3(12.4) | 62.0(14.1) |
| BMI, mean(SD) | 17.3(2.4) | 18.8(3.3) | 17.4(2.5) |
| Parents rating of overall health, mean(SD) | 4.6(0.7) | 4.3(0.6) | 4.6(0.6) |
| History of internalising disorder, Yes(%) | 2(5%) | 2(13%) | 7(37%) |
| History of externalising disorder, Yes(%) | 2(5%) | 8(53%) | 12(63%) |
| Any psychotropic medications, Yes(%) | 0(0%) | 4(27%) | 7(37%) |
| Problems initiating sleep, Yes(%) | 14(34%) | 11(73%) | 16(84%) |
| Problems maintaining sleep, Yes(%) | 10(24%) | 9(60%) | 10(53%) |
| Daytime sleepiness, Yes(%) | 11(27%) | 9(60%) | 11(58%) |
| Any sleep problems, Yes(%) | 22(54%) | 13(87%) | 18(95%) |
| **Home environment:** | | | |
| Household income (AUD), mean(SD) | 87439.0(17787.7) | 86000.0(20546.1) | 74210.5(24903.3) |
| Parent history of mental health disorders, Yes(%) | 15(37%) | 6(40%) | 9(47%) |
| Parent ADHD symptom severity, mean(SD) | 7.4(3.7) | 9.9(4.2) | 11.8(6.2) |
| Angry parenting, mean(SD) | 10.0(2.0) | 12.3(3.5) | 12.7(3.3) |
| **Perinatal:** | | | |
| Maternal smoking during pregnancy, Yes(%) | 5(12%) | 2(13%) | 4(21%) |
| **Pollution:** | | | |
| Air particulate matter (pm) 2.5, mean(SD) | 5.5(0.7) | 5.6(0.5) | 5.2(0.6) |
| **School environment:** | | | |
| Student-teacher total relationship, mean(SD) | 63.0(3.7) | 52.9 (10.1) | 54.8(7.4) |
| **Child stressful life event:** | | | |

| | | | |
|---|---|---|---|
| Relative illness/injury/assault, Yes(%) | 13(32%) | 3(20%) | 8(42%) |
| Death of partner/child's grandparent, Yes(%) | 1(2%) | 1(7%) | 2(11%) |
| Death of close friend/relative, Yes(%) | 17(42%) | 4(27%) | 10(53%) |
| Moved house, Yes(%) | 10(24%) | 5(33%) | 7(37%) |

*BMI=Body Mass Index; QoL=Quality of Life; AUD=Australian currency.*

**Supplementary Table 6.** Results from linear mixed models comparing associations between early life and childhood environment and individual change in DNA methylation by varying ADHD groups (Remitted/Persistent ADHD) to Controls, or by clinical outcomes (Remitted ADHD vs. Persistent ADHD).

| | Probes | Remitted ADHD vs. Controls Estimate (95% CI) | p | Persistent ADHD vs. Controls Estimate (95% CI) | p | Remitted ADHD vs. Persistent ADHD Estimate (95% CI) | p |
|---|---|---|---|---|---|---|---|
| **Child's health:** | | | | | | | |
| QoL: emotional functioning | cg11124426 | 0.04(-0.20, 0.28) | 0.782 | 0.04(-0.23, 0.31) | 0.808 | -0.04(-0.30, 0.23) | 0.812 |
| | cg17657037 | -0.06(-0.30, 0.18) | 0.631 | 0.28(0.01, 0.55) | 0.055 | -0.34(-0.69, 0.01) | 0.094 |
| QoL: school functioning | cg02968116 | -0.26(-0.60, 0.09) | 0.164 | -0.09(-0.32, 0.15) | 0.497 | -0.14(-0.47, 0.18) | 0.432 |
| | cg11124426 | 0.32(-0.06, 0.72) | 0.126 | 0.25(-0.02, 0.51) | 0.087 | 0.11(-0.30, 0.55) | 0.631 |
| | cg17657037 | 0.05(-0.33, 0.44) | 0.792 | 0.28(0.02, 0.54) | 0.051 | -0.24(-0.78, 0.30) | 0.431 |
| QoL: psychosocial health summary score | cg02968116 | -0.10(-0.35, 0.15) | 0.466 | -0.11(-0.35, 0.13) | 0.383 | 0.03(-0.22, 0.26) | 0.848 |
| | cg17657037 | -0.13(-0.40, 0.15) | 0.382 | 0.35(0.09, 0.62) | 0.016 | -0.47(-0.87, -0.08) | 0.037 |
| BMI[a] | cg12881363 | 0.08(-0.19, 0.35) | 0.580 | -0.12(-0.48, 0.24) | 0.547 | 0.21(-0.21, 0.65) | 0.383 |
| | cg17657037 | 0.01(-0.19, 0.20) | 0.943 | 0.02(-0.24, 0.28) | 0.873 | | |
| Parents rating of child's overall health | cg17657037 | 0.01(-0.20, 0.22) | 0.931 | 0.34(0.09, 0.58) | 0.011 | -0.32(-0.67, 0.03) | 0.106 |
| **Home environment:** | | | | | | | |
| Parent ADHD symptom severity | cg17657037 | -0.19(-0.47, 0.08) | 0.205 | -0.15(-0.39, 0.08) | 0.229 | -0.02(-0.38, 0.34) | 0.929 |
| Household income | cg02208776 | -0.27(-0.46, -0.08) | **0.010** | 0.04(-0.14 0.23) | 0.672 | -0.40(-0.62, -0.18) | **0.002** |
| | cg17657037 | -0.14(-0.36, 0.08) | 0.225 | 0.23(0.02, 0.45) | 0.046 | -0.38(-0.72, -0.05) | 0.051 |
| Angry parenting | cg11124426 | -0.14(-0.39, 0.09) | 0.264 | -0.21(-0.47, 0.05) | 0.128 | 0.06(-0.19, 0.29) | 0.673 |
| | cg12881363 | -0.02(-0.35, 0.30) | 0.903 | 0.35(-0.005, 0.69) | 0.069 | -0.37(-0.74, -0.002) | 0.082 |
| | cg17657037 | -0.09(-0.34, 0.15) | 0.474 | -0.03(-0.29, 0.23) | 0.851 | -0.06(-0.39, 0.26) | 0.731 |
| **Pollution:** | | | | | | | |
| Air particulate matter (pm) 2.5 | cg11124426 | 0.07(-0.18, 0.32) | 0.584 | 0.11(-0.15, 0.36) | 0.440 | -0.09(-0.40, 0.22) | 0.620 |
| | cg12577151 | 0.06(-0.07, 0.19) | 0.359 | 0.12(-0.02, 0.25) | 0.111 | -0.06(-0.23, 0.11) | 0.511 |
| **School environment:** | | | | | | | |
| Student teacher relationship – total[a] | cg11124426 | 0.23(-0.04, 0.51) | 0.120 | 0.10(-0.23, 0.43) | 0.582 | 0.12(-0.14, 0.38) | 0.400 |
| | cg12881363 | 0.14(-0.24, 0.52) | 0.499 | -0.13(-0.60, 0.34) | 0.613 | 0.26 (-0.16, 0.69) | 0.273 |
| | cg17657037 | -0.02(-0.30, 0.26) | 0.913 | -0.03(-0.37, 0.31) | 0.882 | | |

*Bold=FDR < 0.05. BMI=Body Mass Index; QoL = Quality of Life. [a]Models were non-converging.*
